# REGEN-COV protects against viral escape in preclinical and human studies

**DOI:** 10.1101/2021.03.10.434834

**Authors:** Richard Copin, Alina Baum, Elzbieta Wloga, Kristen E. Pascal, Stephanie Giordano, Benjamin O. Fulton, Anbo Zhou, Nicole Negron, Kathryn Lanza, Newton Chan, Angel Coppola, Joyce Chiu, Min Ni, Yi Wei, Gurinder S. Atwal, Annabel Romero Hernandez, Kei Saotome, Yi Zhou, Matthew C. Franklin, Andrea T. Hooper, Shane McCarthy, Sara Hamon, Jennifer D. Hamilton, Hilary M. Staples, Kendra Alfson, Ricardo Carrion, Shazia Ali, Thomas Norton, Selin Somersan- Karakaya, Sumathi Sivapalasingam, Gary A. Herman, David M. Weinreich, Leah Lipsich, Neil Stahl, Andrew J. Murphy, George D. Yancopoulos, Christos A. Kyratsous

**Author notes:** These authors contributed equally to this work (Richard Copin, Alina Baum).

## Abstract

Monoclonal antibodies against SARS-CoV-2 are a clinically validated therapeutic option against COVID-19. As rapidly emerging virus mutants are becoming the next major concern in the fight against the global pandemic, it is imperative that these therapeutic treatments provide coverage against circulating variants and do not contribute to development of treatment emergent resistance. To this end, we investigated the sequence diversity of the spike protein and monitored emergence of minor virus variants in SARS-COV-2 isolates found in COVID-19 patients or identified from preclinical in vitro and in vivo studies. This study demonstrates that a combination of non-competing antibodies, REGEN-COV, not only provides full coverage against current variants of concern/interest but also protects against emergence of new such variants and their potential seeding into the population in a clinical setting.

## Introduction

The coronavirus disease 2019 (COVID-19) global pandemic has motivated a widespread effort to understand adaptation of its etiologic agent, the severe acute respiratory syndrome coronavirus 2 (SARS-CoV-2). Similar to all viruses, SARS-CoV-2 is constantly evolving, mutants are selected due to a fitness advantage, which can be associated with an ability to spread more efficiently in the human population or to overcome natural or vaccine-mediated immune responses. Ongoing viral evolution is a major concern for direct antivirals, especially for therapeutics targeting the main proteins recognized by humoral immunity, such as the viral glycoproteins. Such mutations can provide a fitness advantage in the context of drug treatment; therefore they can be rapidly selected during therapy, and may lead to treatment failure and/or spread of treatment resistant-associated variants. To date, multiple studies have investigated the development of viral resistance against neutralizing monoclonal antibodies (mAbs) targeting the SARS-CoV-2 spike protein (Baum et al., 2020a; Liu et al., 2021b; Weisblum et al., 2020)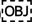. However, better understanding of virus evolution as the virus spreads in humans and the mechanisms driving resistance during treatment is necessary to ensure optimal efficacy of antivirals.

REGEN-COV (previously known as REGN-COV2) is a cocktail of two fully-human non-competing, neutralizing antibodies — casirivimab (REGN10933) and imdevimab (REGN10987) — that target the receptor binding domain (RBD) on the SARS-CoV-2 spike protein and thereby prevent viral entry into the host cell (Hansen et al., 2020). The strategic decision to include a pair of highly potent non-competing antibodies that can simultaneously bind to distinct, non-overlapping epitopes of the RBD was made to safeguard against the rapid emergence of drug resistant variants and provide coverage against circulating viral variants (Baum et al., 2020a). In vitro studies showed that combining two non-competing antibodies protected against rapid escape seen with its individual single-antibody components (Baum et al., 2020a). Additionally, the benefit of this combination approach has been supported by subsequent studies showing that REGEN-COV retains neutralization potency against emerging SARS-CoV-2 variants potentially associated with increased infectivity (e.g. D614G and UK B.1.1.7 variants), as well as those associated with reduced vaccine-induced neutralization titers or vaccine efficacy (e.g. South Africa B.1.351), even when one of its components has partially reduced activity (Garcia-Beltran et al., 2021; Liu et al., 2021a; Madhi et al., 2021; Wang et al., 2021; Yurkovetskiy et al., 2020)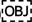. Concerns regarding single antibody approaches in the clinic are emerging, based on increasing prevalence of naturally circulating variants resistant to these single agents(Wang et al., 2021)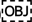, and reports within clinical studies that single antibody treatment is associated with the emergence of resistant (FDA, 2021)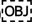.

Although in vitro escape studies using recombinant viruses strongly support the rationale of using non-competing antibody combinations to avoid drug resistance, questions remain regarding the relevance of these findings to authentic SARS-CoV-2 virus, to in vivo infection models, and most importantly, to the clinical setting. In this work we first show that in vitro escape variants are rapidly selected against all individual clinical-stage antibodies targeting the RBD of the spike protein, independent of the targeted epitope or its degree of conservation, that such risk can be mitigated by utilizing combinations of two non-competing antibodies, and even further reduced by combining three non-competing antibodies. We extend these findings by showing that escape variants can also be efficiently selected in vivo upon treatment with individual antibodies, independent of dosage or treatment setting (prophylaxis or therapy), while the REGEN-COV combination fully protects against development of such resistance.

Importantly, we confirm these findings in humans; we investigated the genetic diversity of the entire spike protein across 4,882 samples from 1,000 outpatients or hospitalized patients with confirmed COVID-19 from ongoing Phase 1-3 trials (Weinreich et al., 2021, 2020) (clinicaltrials.gov NCT04666441, NCT04426695, NCT04452318). Analysis of baseline and post-treatment sequence diversity in placebo and REGEN-COV treated patients demonstrated that the non-competing antibody combination protects against selection of drug resistant variants.

## Results

We previously generated a large library of fully human monoclonal antibodies targeting the spike protein of SARS-CoV-2 and selected REGN10933 + REGN10987 as a pair of potently neutralizing molecules that can simultaneously bind to their target (Hansen et al., 2020). We hypothesized that this non-competing antibody combination would retain its potency as the virus evolves in the human population and safeguard against selection of escape mutants following treatment (Baum et al., 2020a; Tegally et al., 2020) With the recent rise in rapidly emerging variants, we are able to test this hypothesis by evaluating the potency of the individual antibodies as well as the REGEN-COV combination against variants of concern/interest as defined by the Centers for Disease Control and Precention (CDC), including B.1.1.7(UK), B.1.427/B.1429 (California), B.1.351 (South Africa), P.1(Brazil), B.1.526 (New York), B.1.617.1/B.1.717.2/B.1.617.3 (India) lineages (CDC, 2021; Davies et al., 2021; Nonaka et al., 2021; Tegally et al., 2020; West et al., 2021; Zhang et al., 2021) Importantly, even in instances where one of the antibodies is impacted, the combination retains full neutralization potency, a key property through which this combination of antibodies was originally selected, thus maintaining coverage against all assessed variants (Fig. 1A&B).

**Figure 1:**
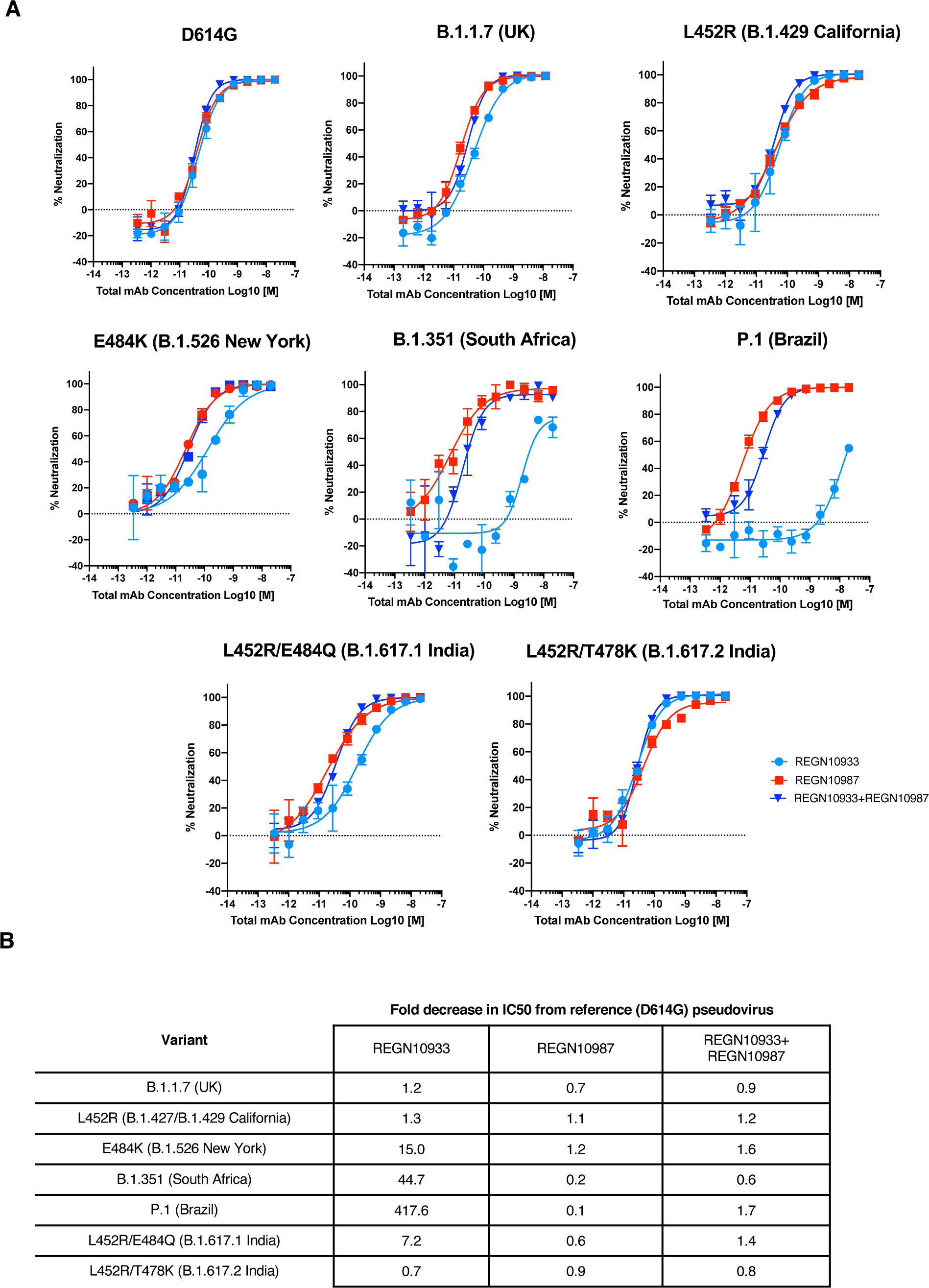
Neutralization of variants of concern and variants of interest by the REGEN-COV combination and individual antibodies. **A)** Neutralization curves and **B)** Neutralization fold-decrease relative to parental D614G variant. Full UK B.1.1.7 (H69del, V70del, Y145del, N501Y, A570D, D614G, P681H, T716I, S982A, D1118H), South African B.1.351 (D80Y, D215Y, L241del, L242del, A243del, L242del, K417N, E484K, N501Y, D614G, A701V) and P.1 (L18F, T20N, P26S, D138Y, R190S, K417T, E484K, N501Y, D614G, H655Y, T1027I, V1176F) variants were assessed. Key RBD residues from B.1.429 California (L452R) and B.1.617.1 India (L452R/E484Q) and B.1.617.2 India (L452R/T478K) variants were assessed. The E484K variant has been identified in many global lineages, including the B.1.526 New York, P.2 Brazil and B.1.618 India lineages.

We have previously shown that combination therapy with two anti-spike antibodies targeting non-overlapping epitopes of the SARS-CoV-2 spike protein minimizes the likelihood of rapid viral escape observed in vitro with monotherapy (Baum et al., 2020), as has also been demonstrated by others (Dong et al., 2021; Weisblum et al., 2020).

Expanding on our previous work using VSV-SARS-CoV-2-spike recombinant virus to monitor resistance to mAb treatment, we now performed such escape studies with several RBD targeting mAbs approved under Emergency Use Authorization (EUA) or in late-stage clinical trials, and tested monotherapy as well as dual and triple combinations. We found that while only one to two passages led to complete virus resistance against all mAbs used as monotherapy, seven consecutive passages were needed to reach complete resistance to the REGEN-COV combination, requiring selection of multiple simultaneous mutations impacting each antibody (Fig.2A&B). Escape variants identified with COV2-2130, COV2-2196, and COV555 antibodies were consistent with those previously described for these molecules (Table S1) (Dong et al., 2021; FDA, 2021)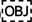. Similar to the REGEN-COV combination, the COV2-2130 and COV2-2196 cocktail was also resistant to rapid escape and full escape was associated with acquiring multiple mutations within the same virus (Fig.2A)(Dong et al., 2021)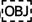. In contrast, the LY-CoV555/CB6 combination behaved similarly to monotherapy, where rapid escape driven by a single amino acid substitution was observed. Such rapid escape from combinations of competing antibodies has been demonstrated in our previous studies and is expected based on the overlapping nature of the LY-CoV555 and CB6 epitopes(Baum et al., 2020a; Jones et al., 2021; Shi et al., 2020; Starr et al., 2021)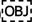.

**Figure 2:**
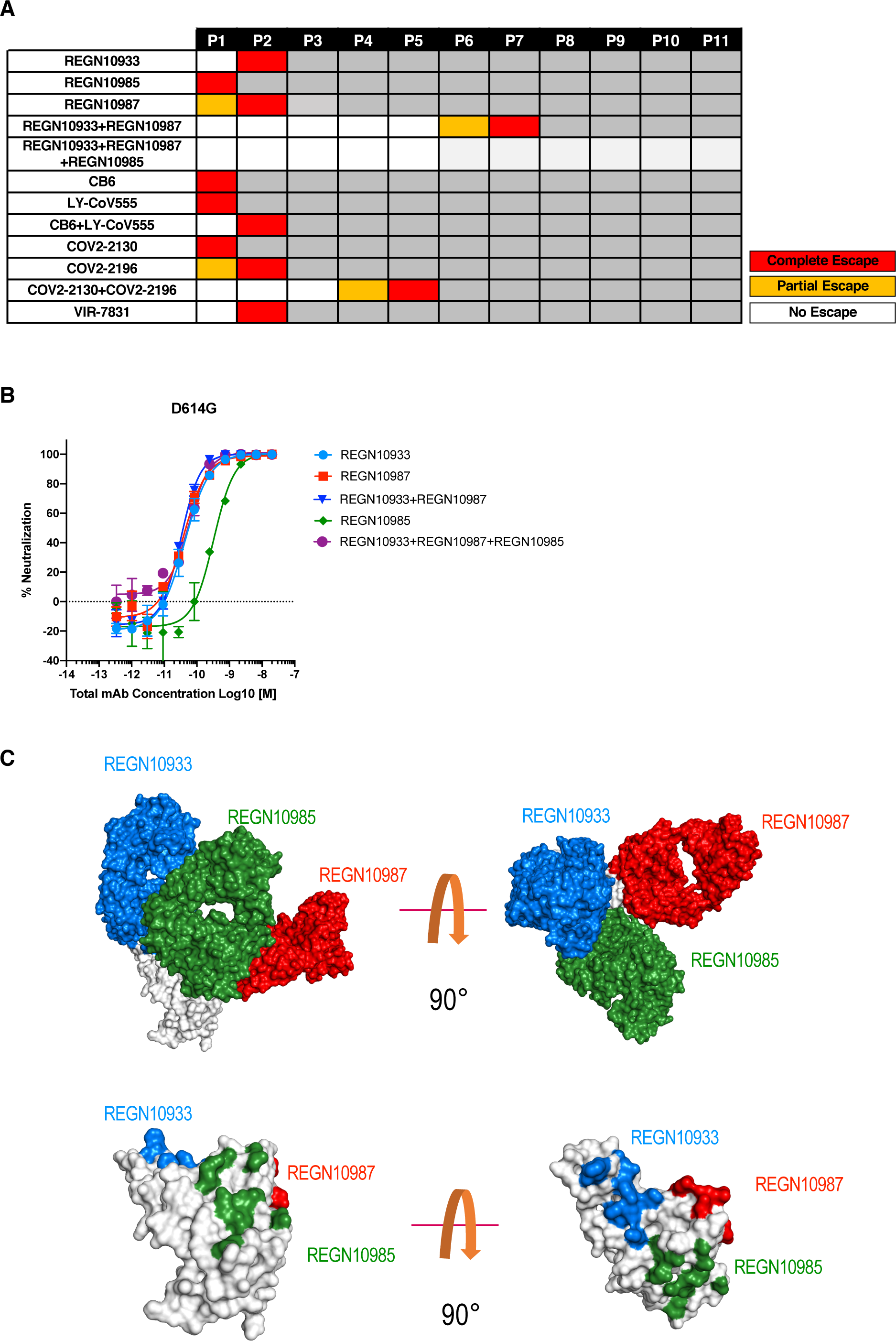
In vitro escape with individual antibodies and antibody combinations. VSV-spike recombinant virus was passaged under pressure of individual antibodies or antibody combinations. **A)** passage number during which the virus completely escaped neutralizing activity of the antibodies as evidenced by the cytopathic effect (CPE), is shown as “complete escape” for wells where antibody concentration was 10ug/ml or above or “partial escape” for wells where antibody concentration was 1-10ug/ml. **B)** Neutralization potency of the two mAb REGN10987+REGN10933 combination and the three mAb REGN10987+REGN10933+REGN10987+REGN10985 combination along with individual mAbs **C)** Two views of the RBD-REGN10933-REGN10987-REGN10985 model. RBD is colored white; REGN10933 is shown blue, REGN10987 is shown in red and REGN10985 is shown in green. Surface representation of the RBD in white with contact patches for each antibody, colors as in panel A.

Importantly, rapid escape observed in the monotherapy setting was independent of the targeted epitope or its sequence conservation as escape variants were rapidly selected against an antibody targeting a broad coronavirus epitope, VIR-7831 (Pinto et al., 2020) (Fig. 2A, Table S1). All VIR-7831 escape residues mapped to the published epitope of the antibody and the resultant escape virus maintained its ability to replicate, indicating that these mutations were not deleterious to the virus (data not shown). Furthermore, the identified escape variants can be found in publicly available genome sequences (GISAID), indicating that they are tolerated by the authentic SARS-CoV-2 virus and thus are likely to be selected under pressure. These experiments highlight the inherent risk of monotherapy against SARS-CoV-2 with any anti-spike mAb, regardless of the targeted epitope or its conservation.

We have previously described a triple antibody combination against Zaire ebolavirus (Pascal et al., 2018) and we similarly decided to assess potential advantages of combining three non-competing neutralizing mAbs targeting the RBD of SARS-CoV- 2:REGN10933+REGN10987+REGN10985. As part of an effort to structurally characterize our SARS-CoV-2 neutralizing antibodies, we determined a cryo-EM structure of REGN10985 (plus another antibody, REGN10989) bound to RBD (Supp. Fig. 4). REGN10985 binds to a broad patch on the side of the RBD, directly below the region contacted by ACE2. Although there is very little overlap between the ACE2 and REGN10985 contact surfaces on the RBD, the REGN10985 Fab is angled so that it overlaps with the space that ACE2 would occupy, making this antibody a direct blocker of ACE2 binding like REGN10933 and REGN10987. By combining this cryoEM structure with our previously-determined structure of REGN10933+REGN10987+RBD (Hansen et al., 2020) we produced a model of the three-antibody complex showing that all three mAbs are able to bind simultaneously to the RBD of the spike protein in a non-overlapping fashion (Figure 2C). We additionally demonstrated that the three-antibody combination has similar neutralization potency as REGEN-COV (Figure 2B). We found that addition of the third non-competing RBD mAb further increased protection against viral escape, with no loss of antiviral potency observed through eleven consecutive passages (Figure 2A, Table S1). These results demonstrate that a three mAb neutralizing non-competing combination targeting the spike RBD provides a potential further advantage as a SARS-CoV-2 therapeutic.

To understand how these in vitro escape findings demonstrating that mAb combination treatments safeguard against development of drug resistance translate to a more physiological setting, we evaluated virus evolution in vivo following mAb administration as both monotherapy (REGN10987 or REGN10933) and as the REGEN-COV combination in the SARS-CoV-2 hamster challenge model. Animals were challenged with SARS-CoV-2 (USA-WA1/2020) and treated with either high or low dose (50mg/kg or 5mg/kg) of either REGN10987 or REGN10933 as monotherapy or with the REGEN-COV combination (50mg/kg or 5mg/kg) in both prophylactic and treatment settings (Figure 3A). These doses were previously shown to provide partial or complete efficacy in this animal model with REGEN-COV (Baum et al., 2020b). Lungs from infected animals were collected on day 7 post-challenge, presumably after the virus has had an opportunity to go through multiple rounds of replication, to isolate viral RNA. RNAseq analysis was used to detect variants in the spike protein present even at low frequencies in the virus population (>5%). To identify putative treatment resistant variants, we focused our analysis on the spike RBD, where the REGEN-COV antibodies are known to bind, and where all previously identified resistant variants to either antibody are located. Analysis of the RBD sequence in saline treated animals did not reveal any variants, indicating that in the absence of treatment the virus is not under strong selection pressure in this model (Fig. 3B&C). However, several variants in the RBD were identified in animals treated with either of the single antibodies as monotherapy (Fig. 3B&C). Remarkably, our analysis demonstrated selection of resistance variants in almost half (18/40) of monotherapy treated animals versus none (0/20) of the animals treated with the REGEN-COV combination. Out of the 7 RBD variants identified in monotherapy treated animals, 6 have been previously described as resistance variants or mapped to the same amino acid position as other resistance variants to either REGN10933 or REGN10987 (Table S3). Four of these variants were previously identified in our in vitro escape studies, validating the relevance of using VSV-SARS-CoV-2-spike recombinant virus to identify SARS-CoV-2 escape mutations (Baum et al., 2020a). Interestingly, variant selection occurred with monotherapy treatment in both the prophylactic and therapeutic settings, and in both low and high dose groups. Although rare unique variants outside of the RBD were also identified in a single animal treated with the REGEN-COV combination (Fig. 3B&C), the low frequency of these variants and their location well outside the epitopes of the antibodies make it extremely unlikely that they were associated with the REGEN-COV treatment.

**Figure 3:**
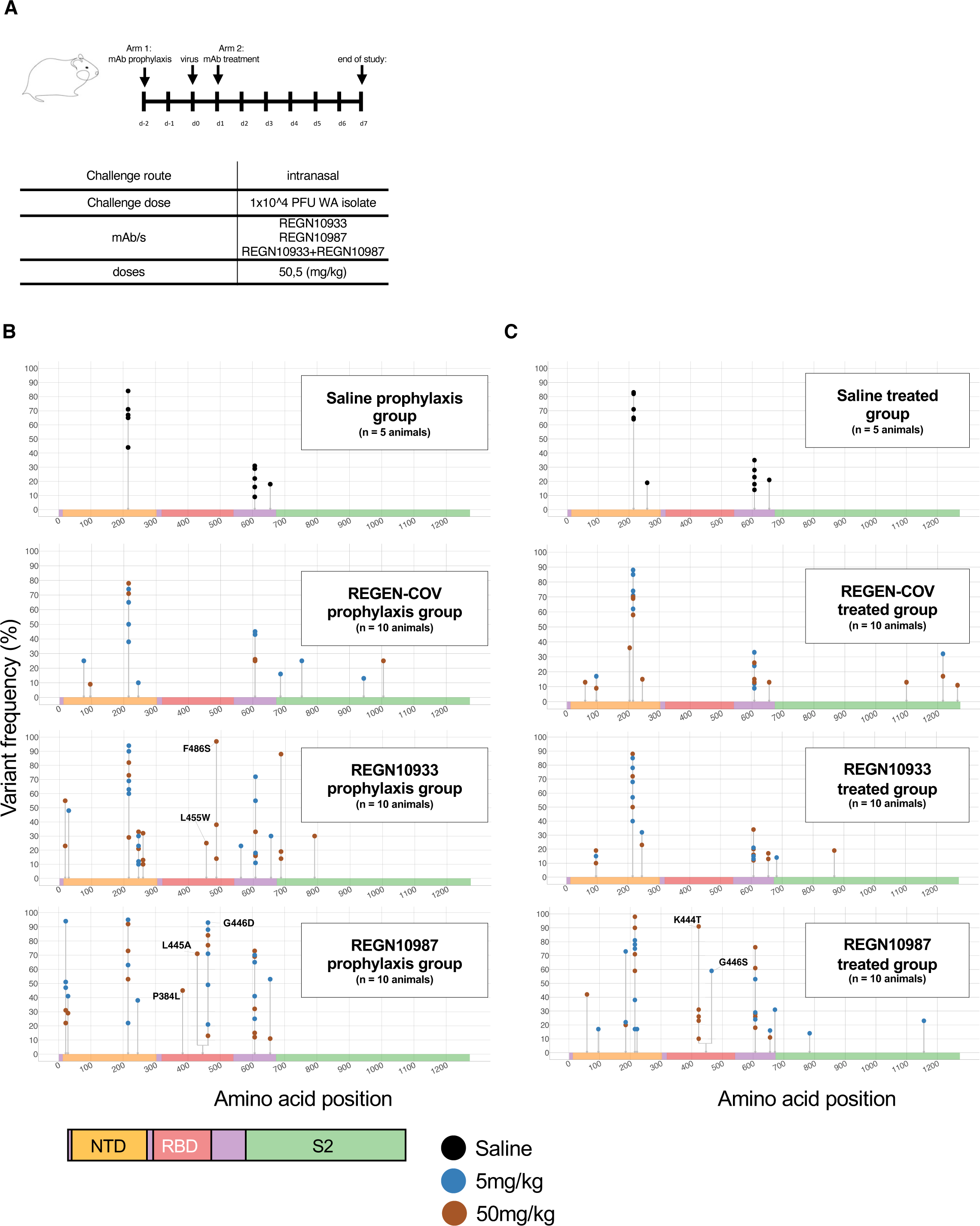
Frequency of SARS-CoV-2 spike variants identified in viruses from lungs of hamsters treated with either REGN10987 or REGN10933 as monotherapy or with the REGEN-COV combination in both prophylactic and treatment settings. **A)** Study design. **B,C)** Graphs show the distribution and frequencies of amino acid variants across the spike protein sequence identified in samples from the **B)** prophylactic or **C)** treated groups. Mutated sites are indicated with grey lines and dots show the frequency of variants in each animal. The color of each dot indicated the dose used. Spike protein domain are color coded as indicated and all amino acid changes in RBD are labeled.

To assess the emergence of drug resistance in REGEN-COV treated COVID-19 patients, we characterized the genetic diversity of 1,336 and 3,546 SARS-CoV-2 RNA samples isolated from 272 hospitalized and 728 non-hospitalized individuals, respectively, at baseline and several time points post mAb cocktail administration (Table S2). In both hospitalized and non-hospitalized study groups, patients were randomly assigned (1:1:1) to receive placebo, 2.4g of REGEN-COV, or 8.0g of REGEN-COV, and patient samples were collected at multiple time points from baseline (day 1) to study end point (day 29). RNA-seq analysis was performed to characterize the genetic diversity of the spike sequence from all swabs positive for SARS-CoV-2 RNA by qRT-PCR. Minor variants present at a frequency of at least 15% in the virus population were identified. In total, 244 and 514 amino acid changes were identified across all samples from hospitalized and non-hospitalized patients, respectively. For both studies, variant counts and frequencies in all three treatment groups were similar, indicating that REGEN-COV did not increase spike protein diversity relative to placebo (Table S2).

To determine whether REGEN-COV treatment contributed to selection of resistant variants in the spike protein, we longitudinally compared the frequency of minor variants in each patient from all available RT-PCR positive samples. Among all patient samples from the hospitalized group, we identified a total of 89, 87 and 108 amino acid changes in the placebo, low-dose and high dose treated groups, respectively. These mutations were evenly spread across the entire spike protein sequence (Fig. 4A&B). The low number of variants in the RBD found among samples from patients treated with REGEN-COV further demonstrated lack of treatment emergent selection relative to placebo (15 RBD variants in placebo versus 12 low-dose/12 high-dose in REGEN-COV treated group). Only 17% (42/244) of all amino acid changes were observed at more than one time point in at least one patient (Fig. 4B) indicating that the vast majority of variants were lost over time likely as a result of lack of fitness or population bottlenecks. Three of these variants were found in the RBD in the REGEN-COV treated groups only (S477N, N501T and K537R). The S477N variant was present in 7 REGEN-COV treated patients at multiple time points, including baseline (Fig. S1). S477N is a common SARS-CoV-2 variant present at a frequency of >5% viruses circulating worldwide (source: GISAID). K537R was identified in 1 treated patient and is a rare mutation, which has been found in only <10 virus isolates worldwide to date (Fig. 5, source: GISAID). N501T was observed in 3 treated patients in this study (Fig. 5). Of note, in all patients, the three variants were identified at baseline or earliest time points (<5 days after REGEN-COV treatment) and no significant increase in their frequencies between early and late time points were observed indicating absence of treatment related selection.

**Figure 4:**
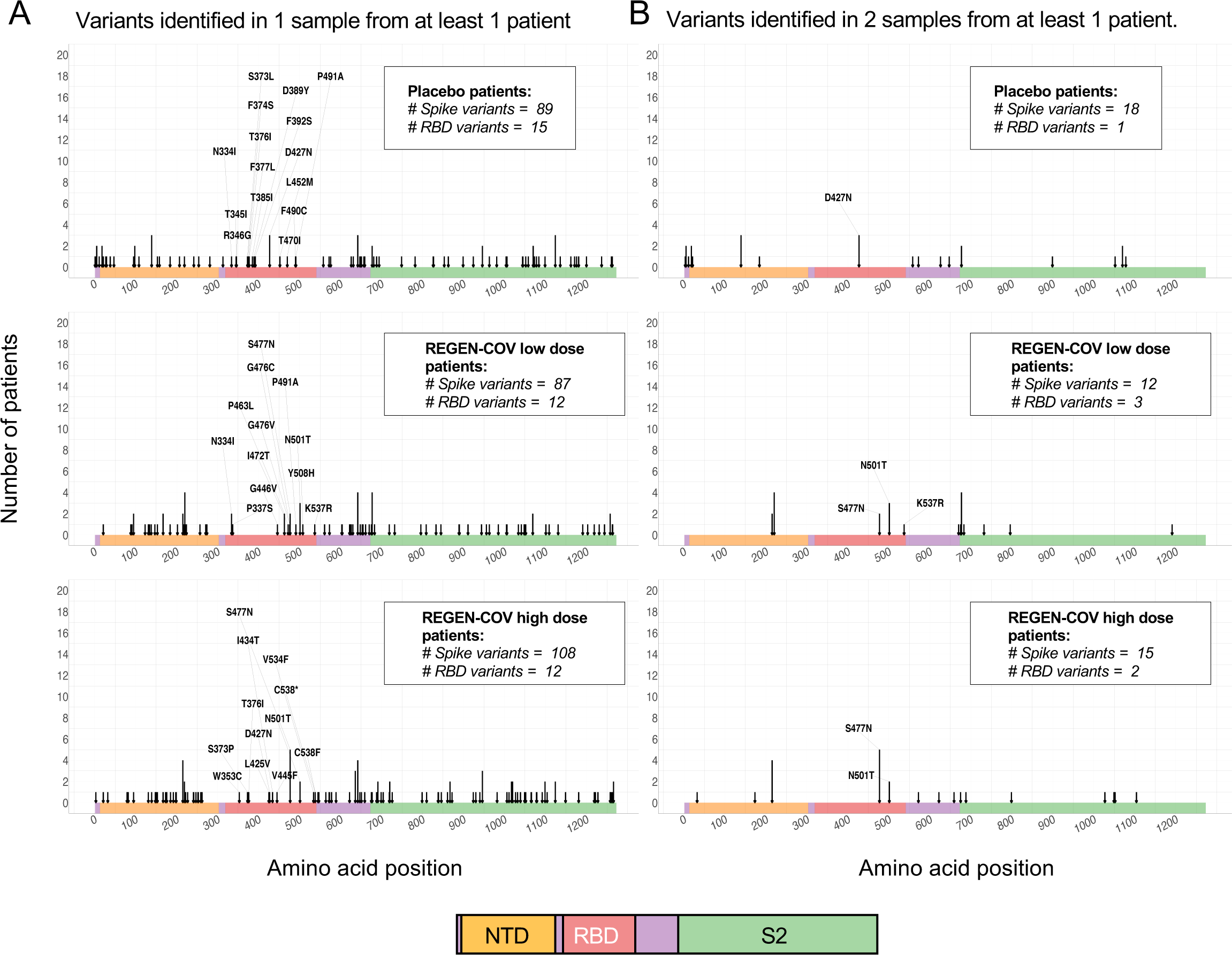
Variant analysis of virus sequences in SARS-CoV-2 positive samples from hospitalized patients. Graphs show the distribution of amino acid variants across the spike protein sequence identified in samples from the placebo, REGEN-COVlow- dose and REGEN-COVhigh-dose treated patients. Mutated sites are indicated with arrows and arrow length designate the number of patients with 1 or more variant- containing samples. Spike protein domains are color coded as indicated and all amino acid changes in RBD are labeled. **A)** Arrows highlight variants identified in 1 or more samples from at least one patient. **B)** Arrows highlight variants identified in 2 or more samples from at least one patient. All samples were characterized by the defining Glycine substitution at position 614.

**Figure 5:**
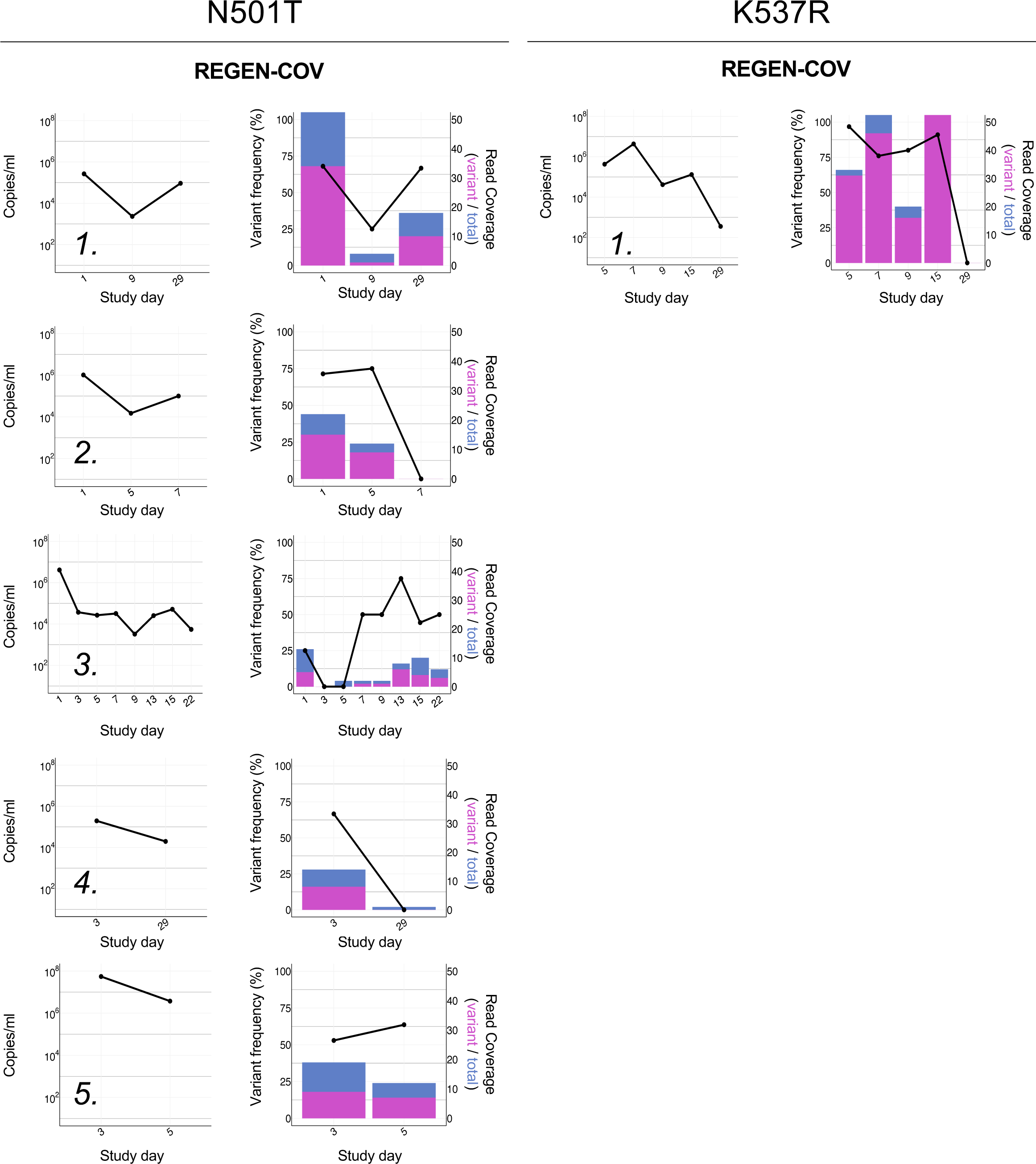
Viral load and longitudinal frequencies of variants in REGN-COV2- treated, SARS-CoV-2 positive samples from hospitalized patients. Viral load was determined by quantitative RT-PCR. **A)** N501T and **B)** K537R variant frequencies are indicated by the black line as percent of reads with nSNPs relative to total number of covering reads. Histograms show total (blue) and mutated (magenta) read coverage at variant position.

We identified a total of 216, 248 and 212 amino acid changes among the 1,215 placebo, 1,203 low-dose and 1,128 high-dose REGEN-COV treated patient samples respectively from the non-hospitalized group, spread across the entire spike protein sequence (Fig. 6). The RBD contained respectively, 23, 27 and 31 amino acid changes among which S477N and only 2 other variants (S494P and G446V) were identified at more than one time point in at least one REGEN-COV treated patient (Fig. 6B). The S477N variant was present in 3 REGEN-COV and 5 placebo treated patients at multiple time points, including baseline (Figure S2). S494P is a rare circulating mutation (∼0.15% GIDAID) only found at multiple time points (including baseline) in 2 REGEN-COV treated patients. G446V is a rare variant (∼0.02% GISAID) identified in 4 REGEN-COV treated and 2 placebo patients (Fig. 7). One of these patients (Fig. 7 - Patient B4), represented the only example with a consistent late increase in the relative frequency of the variant, however, it should be pointed out that there is no evidence that the absolute frequency of the variant increased over time as viral levels dropped dramatically over time. For all G446V patients, including Patient B4, we observed that the frequency of variants was decreased to undetectable at the latest time points, indicating that drug mediated selection is unlikely.

**Figure 6:**
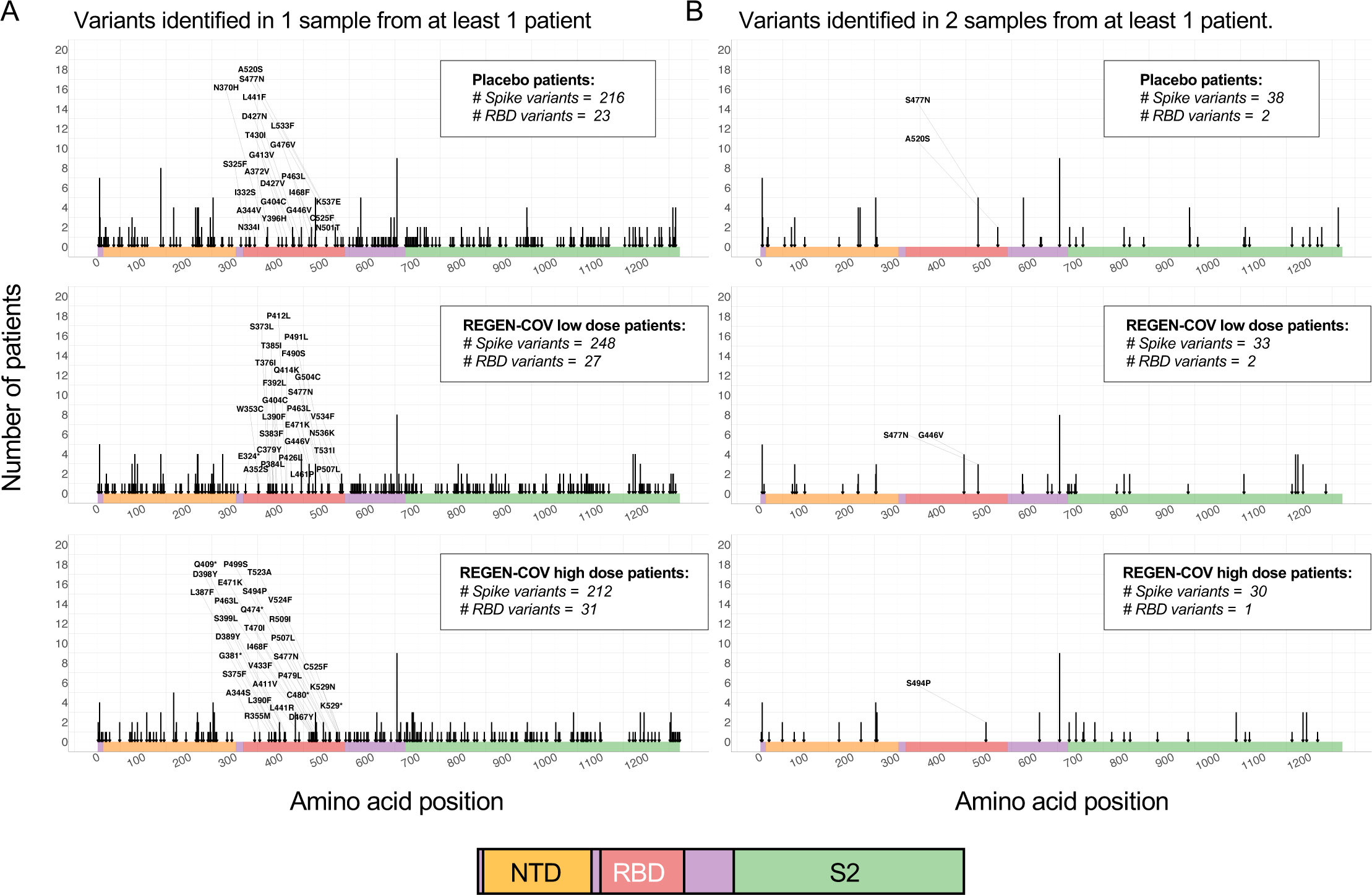
Variant analysis of virus sequences in SARS-CoV-2 positive samples from non-hospitalized patients. Graphs show the distribution of amino acid variants across the spike protein sequence identified in samples from the placebo, REGEN-COV low-dose and REGEN-COV high-dose treated patients. Mutated sites are indicated with arrows and arrow length designate the number of patients with 1 or more variant- containing samples. Spike protein domains are color coded as indicated and all amino acid changes in RBD are labeled. **A)** Arrows highlight variants identified in 1 or more samples from at least one patient. **B)** Arrows highlight variants identified in 2 or more samples from at least one patient. All samples were characterized by the defining Glycine substitution at position 614.

**Figure 7:**
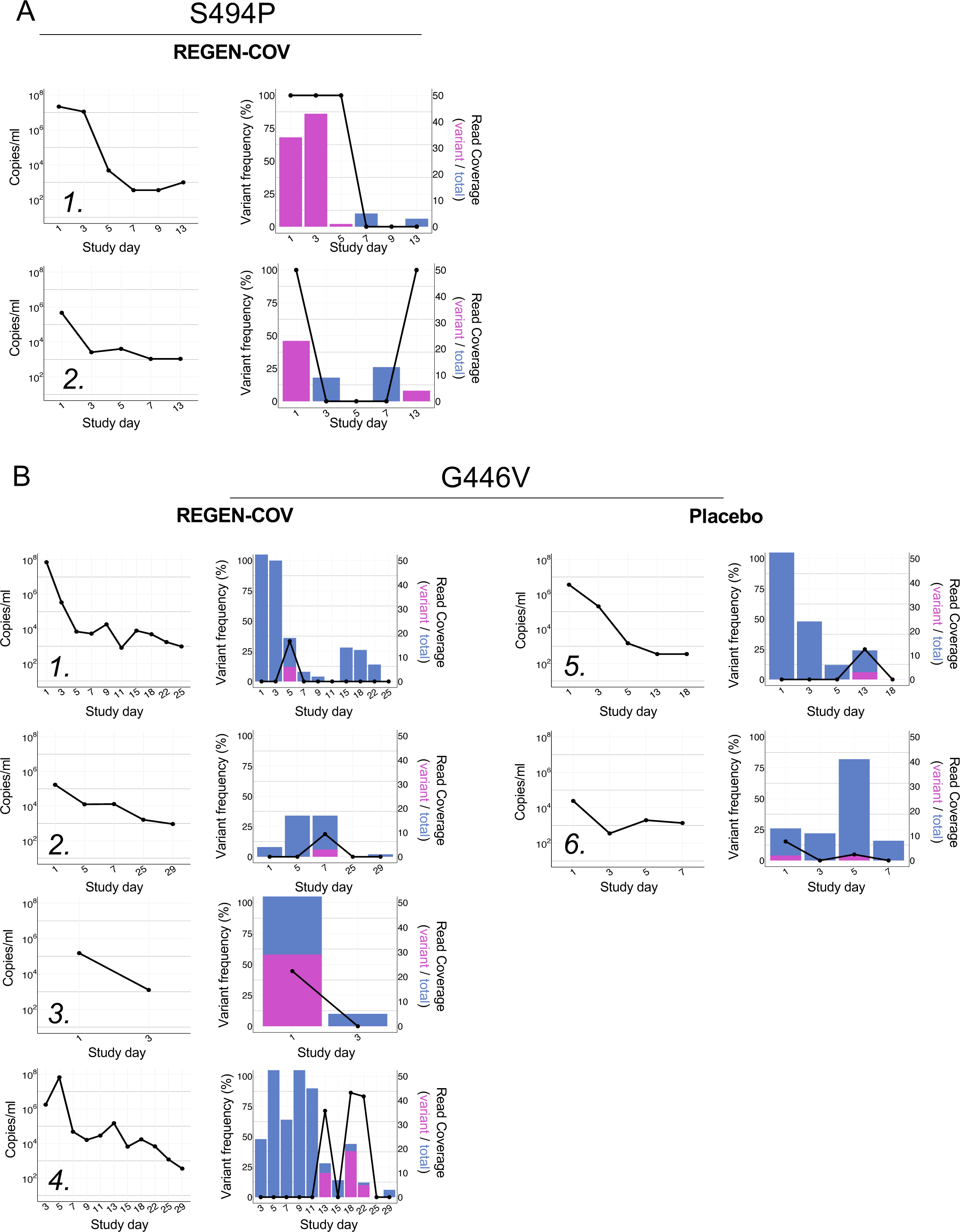
Viral load and longitudinal frequencies of variants in SARS-CoV-2 positive samples from non-hospitalized patients. Viral load was determined by quantitative RT-PCR. **A)** S494P, **B)** G446V and variant frequencies are indicated by the black line as percent of reads with nSNPs relative to total number of covering reads. Histograms show total (blue) and mutated (magenta) read coverage at variant position.

To determine the functional impact of RBD variants identified at multiple time points in REGEN-COV treated patients, individual antibodies and the combination were assessed for neutralization potency against the G446V, S494P, S477N, and K537R variants (Fig. S3). Decrease in neutralization potency was observed with REGN10987 (135-fold) against the G446V variant, and minimally with REGN10933 (4.5-fold) against the S494P variant, whereas no impact was observed on either mAb with the S477N or K537R variants. The N501T variant was not assessed but an alternative substitution N501Y did not show any impact on either antibody or the REGEN-COV combination.

Importantly, in all instances the REGEN-COV combination retained full neutralization potency, thereby providing its full antiviral activity in treated individuals and limiting any potential selection of resistant variants.

## Discussion

The increasing use of monoclonal antibodies as a clinically validated therapeutic option against SARS-CoV-2 raises questions about which of these therapies may be impacted by circulating and rapidly emerging variants and which may induce treatment emergent resistance. The recent recognition that some immunodominant residues in the spike protein, such as E484K, often impact both mAb therapies and vaccine-induced immune responses makes it imperative to ensure that mAb treatments not only provide protection against common circulating variants, but to also ensure that treatment-induced escape variants are not seeded into the population, where they may contribute to undermining efficacy of other therapeutics or vaccines (Greaney et al., 2021; Wang et al., 2021). Real-world manifestations of such concerns are demonstrated by the recent withdrawal of emergency use authorization of bamlanivimab monotherapy due to high prevalence of the B.1.526 and B.1.427/B.1429 variants, as well as the recent US government recommendation against the use of the bamlanivimab/estevimab combination in Illinois and Massachusetts due to high prevalence of the P.1 variant, which impacts both antibodies in the combination(HHS, 2021b, 2021a).

To date, we and others have identified antibody resistant variants through in vitro escape studies and have demonstrated that combinations of two non-competing antibodies provide protection against rapid viral escape (Baum et al., 2020a; Weisblum et al., 2020). We hypothesized that a cocktail of three non-competing antibodies could further delay emergence of resistance. Although, neutralizing antibodies against the N-terminal domain (NTD) of spike have been described (Liu et al., 2020), this region is frequently deleted in emerging variants (Wang et al., 2021), therefore such molecules are not ideal as therapeutics. We thus screened for and identified a third, non-competing, RBD-binding mAb and now show that a triple antibody combination containing this mAb is even less susceptible to escape than the double combination.

However, the question regarding the relevance of these in vitro escape studies remains, and it is important to examine viral dynamics under therapeutic antibody pressure in more relevant settings, including in in vivo efficacy models and in treated patients. Our study, for the first time, provides evidence that in vitro escape studies with recombinant virus are highly predictive of in vivo selection. Indeed, we are able to show that in animals and in humans, the relative risk of selecting for spike variants that are resistant to treatment is predicted by in vitro findings. Treatment with the single antibody components of the REGEN-COV combination results in rapid selection of escape variants, whereas the combination protects against viral escape in vitro, in vivo, and in COVID-19 patients. Furthermore, by comparing escape mutations identified through in vitro assays utilizing recombinant VSV-based virus to those identified in SARS-CoV-2 infected hamsters, we show that many identical mutations are identified in both settings, further validating the in vitro findings.

Understanding pathogen population diversity and selection within the infected host has important implications for understanding virus evolution and resistance processes, and for proper identification of appropriate drug treatments. To date, the vast majority of SARS-CoV-2 sequencing data has come from global surveillance efforts and reporting of consensus sequences that include fixed mutations in isolates obtained from independent individuals. This is the first large-scale report describing, within individual patients, the longitudinal evolution of the SARS-CoV-2 virus population. Moreover, this study is the first to evaluate the effects of antiviral antibody treatment on virus evolution in humans. As predicted by our in vitro and in vivo preclinical studies, treatment with the REGEN-COV cocktail in the clinic did not seemingly select for resistant variants nor appreciably alter virus evolution. Our approach allowed us to observe minor variants and our data indicate the vast majority of infected individuals are already harboring spike protein variants at baseline, prior to use of any antiviral agent and before natural immunity is established, as half of the study participants had not yet mounted an antiviral immune response at baseline (Weinreich et al., 2020). Indeed, from our clinical sequencing data we can observe multiple variants in amino acid residues identified in our in vitro escape studies expected to impact clinical antibody candidates, including VIR-7831 (P337, T345) and COV-2196 (G476) as well as our own individual antibodies. This clearly demonstrates that the SARS-CoV-2 circulating population is highly heterogeneous in infected patients and the large reservoir of existing (or future) minor variants could be readily selected under the right selective pressure conditions. Indeed, treatment emergent resistant variants have been reported in clinical studies in which patients were treated with anti-SARS-CoV-2 antibody monotherapy (FDA, 2021).

Although the REGEN-COV combination maintains its neutralization potency against all variants of interest/concern to date, one of its components (REGN10933) demonstrates partially decreased potency against some variants. This highlights the potential risk that even non-competing combination therapies may not retain their full advantages in some geographical regions with high prevalence of select variants of concern. Thus, active surveillance of circulating viruses and development of next-generation antibody combinations where multiple components maintain coverage against emerging variants should be an important focus for the future of antiviral mAb therapeutics. This type of approach will require agility and may need to be analogous to the framework applied to seasonal updates of influenza vaccine components, where new anti-spike mAbs are substituted or added as necessary to maintain optimal efficacy against a rapidly mutating pathogen.

To ensure optimal and sustained efficacy, an antiviral therapeutic intervention should be prospectively designed to not only have coverage against major circulating variants, but also to prevent the selection of minor escape variants. And although the risk of viral resistance cannot be completely eliminated, it can be greatly reduced through selection of optimal combination therapies. This study demonstrates that carefully designed antibody combinations can achieve both of these critical objectives.

## Limitations of the Study

The current study focused on assessing likelihood of identifying REGEN-COV mediated selection of resistant viral variants in treated COVID-19 patients and overall predictability of preclinical in vitro and in vivo escape studies to human patients. Our data clearly demonstrated that treatment with any RBD-directed mAb monotherapy results in rapid selection of drug resistant viral variants. However, although we did not identify any clear selection of resistant variants in the clinical population after treatment with REGEN-COV, it was impossible for us to directly compare monotherapy and combination therapy in the clinical setting due to the combination nature of our therapeutic. Thorough sequencing of viral samples post-treatment with clinical monotherapy products may help to provide clinical data on likelihood of resistance in that setting. Furthermore, although we assessed over 3000 samples from 1000 patients in our study to identify potential REGEN-COV associated selection of variants it is possible that this sample size may not be large enough to identify very rare events.

## Acknowledgements

The authors would like to thank Kristen Tramaglini, Yasmin Khan, Lisa Boersma and Sarah Whittleton for their continuous support.

## Author Contributions

A.B. and C.A.K. conceptualized the studies. E. W., K.E.P., S.G., and B.O.F. conducted escape studies and neutralization assays. A.Z., N.N., K.L., N.C., A.C., J.C., M.N., Y.W., and G.S.A. performed RNA sequencing and data analysis. H.M.S., K.A., and R.C. carried out the hamster experiments. A.R.H., K.S., Y.Z., and M.C.F. performed structural analysis. A.T.H., J.D.H., S.A., T.N., S.S-K., S.S., G.A.H., D.M.W planned and carried out clinical studies. A.T.H., J.D.H. oversaw clinical sample and data acquisition. S.M., and S.H. oversaw clinical data acquisition and data analysis. L.L. oversaw program direction. R.C., A.B., N.S., A.J.M, G.D.Y, C.A.K analyzed data. R.C., A.B., G.D.Y., C.A.K. wrote the manuscript.

## Methods

### RESOURCE AVAILABILITY

#### Lead contact

Further information and requests for resources and reagents should be directed to and will be fulfilled by the Lead Contact, Christos Kyratsous.

#### Materials availability

Regeneron materials described in this manuscript may be made available to qualified, academic, noncommercial researchers through a materials transfer agreement upon request at https://regeneron.envisionpharma.com/vt_regeneron/. For questions about how Regeneron shares materials, use the email address.

#### Data and code availability

All relevant data are available from the authors upon request. Protein Data Bank (PDB) identification number for the SARS-CoV-2 RBD complex with REGN10985 and REGN10989 Fabs is 7M42. The corresponding cryo-EM map is in the Electron Microscopy Data Bank (EMDB) with identification number EMD-23662.

### EXPERIMENTAL MODEL AND SUBJECT DETAILS

#### Human Subjects

The outpatient study is an ongoing adaptive, multicenter, randomized, double-blind, placebo-controlled, seamless phase 1/2/3 clinical trial in COVID-19 outpatients (clinicaltrials.gov: NCT04425629). Non-hospitalized patients > 18 years of age with a SARS-CoV2-positive test result <72 hours, and symptom onset < 10 days before randomization, were randomized 1:1:1 to receive placebo, 2.4g (1.2 g each of casirivimab and imdevimab) or 8.0 g (4.0 g each of casirivimab and imdevimab).

The hospitalized study is an ongoing adaptive, multicenter, randomized, double-blind, placebo-controlled, seamless phase 1/2/3 clinical trial in hospitalized patients with Covid-19 (clinicaltrials.gov: NCT04426695). Hospitalized patients > 18 years of age with a SARS-CoV2-positive test result <72 hours, and symptom onset < 7 days before randomization, were randomized 1:1:1 to receive placebo, 2.4g (1.2 g each of casirivimab and imdevimab) or 8.0 g (4.0 g each of casirivimab and imdevimab).

#### Animals

A total of 80 golden hamsters, male, approximately 9 weeks of age (130-150g) were used in the study. Animals were housed one to five per cage; caging complies with the cage requirements (type, floor space, height, etc.) of the Animal Welfare Act and the Guide for the Care and Use of Laboratory Animals. Animals were acclimated to ABSL-3 laboratory conditions prior to exposure. In the ABSL-3, prior to virus exposure, animals were observed at least once daily, and after virus exposure animals will be observed at least twice daily for morbidity and mortality. On Day -2 relative to exposure, animals were sedated and received treatment via intraperitoneal injection.

#### Cells

African green monkey kidney cells (Vero, ATCC CCL-81), Vero clone E6 (Vero E6, ATCC CRL-1586), and human embryonic kidney cells (293T, ATCC CRL-3216), were maintained at 37°C and 5% CO2 and cultured in high glucose Dulbecco’s Modified Eagle’s Medium without glutamine (DMEM; Life Technologies) supplemented with 10% fetal bovine serum (FBS; Life Technologies) and 1% Penicillin/Streptomycin/L-glutamine (Life Technologies).

#### Viruses

SARS-CoV-2 generated from isolate USA-WA1/2020 was used for animal exposures. A fourth cell-culture passage (P4) of SARS-CoV-2 was obtained from Biodefense and Emerging Infections Research Resources Repository (BEI Resources, catalog number NR-52281, GenBank accession number MN985325.1) and propagated at Texas Biomed. The stock virus was passaged for a fifth time in Vero E6 cells at a multiplicity of infection (MOI) of approximately 0.001. This master stock was used to generate a sixth cell culture passage exposure stock by infecting VeroE6 cells at an MOI of 0.02. The resulting stock had a titer of 1.2 x 10^6^ PFU/mL. The exposure stock has been confirmed to be SARS-CoV-2 by deep sequencing and was identical to the published sequence (MN985325.1) at the consensus level.

### METHOD DETAILS

#### Generation of recombinant VSV

Non-replicative pseudoparticles were generated using a VSV genome encoding the firefly luciferase and GFP genes instead of the native viral glycoprotein (VSV-G).

Infectious particles complemented with VSV-G (VSV-ΔG-Fluc-2A-GFP/VSV-G) were recovered and produced using standard techniques with minor modifications (Baum et al., 2020a; Hansen et al., 2020; Lawson et al., 1995). HEK293T cells (ATCC CRL-3216) were plated on poly-lysine treated plates and incubated overnight in DMEM without glutamine (Life Technologies), 10% fetal bovine serum (Life Technologies) and 1% Penicillin/Streptomycin/L-glutamine (Life Technologies). The following day, the cells were transfected with the VSV genomic clone driven by a T7 promoter and helper plasmids expressing the VSV-N, VSV-P, VSV-G, VSV-L, and T7 RNA polymerase with Lipofectamine LTX reagent (Life Technologies). After 48 hours, the transfected cells were co-cultured with BHK-21 cells (ATCC CCL-10) transfected with VSV-G using the SE cell Line 4D-Nucleofector X Kit L (Lonza) in DMEM without glutamine (Life Technologies), 3% fetal bovine serum (Life Technologies) and 1% Penicillin/Streptomycin/L-glutamine (Life Technologies). Cells were monitored for GFP expression or cytopathic effect (CPE) indicative of virus replication. Virus was then plaque purified, expanded, and titered in BHK-21 cells transiently expressing VSV-G. Fully replicative VSV-SARS-CoV-2-S virus was generated by replacing the VSV glycoprotein with the native SARS-CoV-2 sequences encoding residues 1-1255 of the spike protein (MN908947.3). VSV-SARS-CoV-2-Spike virus was recovered as described above but the HEK293T cells were instead cocultured with BHK-21 cells transfected with both VSV-G and hACE2. VSV-SARS-CoV-2-S virus was plaque purified and titered in Vero cells (ATCC CCL-81) and expanded in Vero E6 cells (ATCC CRL-1586). After collection, stocks of both viruses were centrifuged at 3000xg for 5 minutes to clarify, sucrose cushioned to concentrate 10-fold, aliquoted, and frozen at - 80C.

#### Pseudotyping of VSV

Non-replicative pseudoparticles were generated as previously described (Baum et al 2020). Human codon-optimized SARS-CoV-2 spike (MN908947.3) was cloned into an expression plasmid. A total of 1.2 x 107 HEK293T cells (ATCC CRL-3216) were seeded overnight in 15-cm dishes in DMEM without glutamine (Life Technologies) containing 10% heat-inactivated fetal bovine serum (Life Technologies), and Penicillin-Streptomycin-L-Glutamine (Life Technologies). The following day, the cells were transfected with 15ug spike expression plasmid with Lipofectamine LTX (Life Technologies) following the manufacturer’s protocol. At 24 hours post transfection, the cells were washed with phosphate buffered saline (PBS) and infected at a MOI of 1 with the VSV-ΔG-Fluc-2A-GFP/VSV-G virus diluted in 10mL Opti-MEM (Life Technologies). The cells were incubated 1 hour at 37C with 5% CO2. Cells were washed three times with PBS to remove residual input virus and overlaid with DMEM with glutamine (Life Technologies) with 0.7% IgG-free BSA (Sigma), sodium pyruvate (Life Technologies), and Gentamicin (Life Technologies). After 24 hours at 37C with 5% CO2, the supernatant containing pseudoparticles was collected, centrifuged at 3000xg for 5 minutes to clarify, aliquoted, and frozen at -80C. Variants were cloned into the spike expression plasmid using site-directed mutagenesis and pseudoparticles were produced as described above.

#### Neutralization assays with VSV based pseudoparticles

Vero cells (ATCC: CCL-81) were seeded in 96-well black, clear bottom tissue culture treated plated (Corning: 3904) at 20,000 cells/well in DMEM media without glutamine (Life Technologies) containing 10% heat-inactivated fetal bovine serum (Life Technologies), and 1X Penicillin/Streptomycin/L-Glutamine (Life Technologies) 24 hours prior to assay. Cells were allowed to reach approximately 85% confluence before use in assay. Antibodies were diluted in Infection Media containing DMEM with glutamine (Life Technologies), 0.7% Low IgG BSA (Sigma), 1X Sodium Pyruvate (Life Technologies), and 0.5% Gentamicin (Life Technologies) to 2X assay concentration and diluted 3-fold down in Infection media, for an 11-point dilution curve in the assay beginning at 3 ug/mL (20 nM). Antibody dilutions were mixed 1:1 with pseudoparticles for 30 minutes at room temperature prior to addition onto Vero cells. Cells were incubated at 37C, 5% CO2 for 24 hours. Supernatant was removed from cells prior to lysis with 100 uL Glo Lysis Buffer (Promega). 100 uL resuspended Bright Glo substrate (Promega) was then added and luminescence was read on a Spectramax i3x (Molecular Devices). Exported values were analyzed using GraphPad Prism (v8.4.1)

#### In vitro Escape Studies

Escape studies were performed with recombinant VSV-SARS-CoV-2-Spike virus as previously described (Baum et al., 2020a). Escape variants were selected by incubating VSV-SARS-CoV-2-Spike under antibody pressure ranging from 0.016ug/mL to 50ug/mL. After 30 minutes of incubation, the mixture was used to infect 2.5x10^5 Vero (ATCC: CCL-81) or 3x10^5 Vero E6 (ATCC CRL-1586) cells at a multiplicity of infection (MOI) of 5. Virus replication was monitored by screening for cytopathic effect over 96 hours. The supernatant and cellular RNA was collected from the wells with the highest antibody concentration with evident virus replication. The RNA was then subjected to RNAseq to identify spike protein variants. For subsequent rounds of selection,100uL of supernatant containing the virus was passaged under the same or greater antibody concentrations as in previous passages until complete CPE was observed after antibody treatment at concentration of ≥10 ug/mL.

#### Hamster Study

A total of 80 golden hamsters, male, approximately 9 weeks of age (130-150g) were used in the study. Animals were weighed prior to the start of the study. The animals were monitored twice daily for signs of COVID-19 disease (ruffled fur, hunched posture, responsiveness, nasal discharge, labored breathing, a.o.) during the study period. Body weights were measured once daily during the study period. Antibodies were dosed through intraperitoneal (IP) injection (0.200ml volume). Animals were challenged with 1x10^4 PFU of (USA-WA1/2020 (NR-52281; BEI Resources) by administration of 0.05ml total of viral inoculum, 0.025ml dropwise into each nostril (mock exposed animals received the same volume of sterile PBS). Prior to treatment and virus administration, animals were sedated using inhalational isoflurane. At study end (day 7 post-challenge), animals were euthanized via inhalational CO_2_ and tissues samples were aseptically collected for RNAseq. The day 7 time-point was selected for RNA collection and analysis as the virus has had ample ability to replicate in the animal allowing for detection of selected variants. Tissue samples were inactivated using Trizol Reagent (Invitrogen, Carlsbad, CA, USA). All animal studies were conducted in compliance with all relevant local, state, and federal regulations and were approved by the Texas Biomed Institutional Animal Care and Use Committee (IACUC; protocol number 1739MA).

#### Clinical Trial Design

##### Outpatient clinical trial

This is an ongoing adaptive, multicenter, randomized, double-blind, placebo-controlled, seamless phase 1/2/3 clinical trial in COVID-19 outpatients (clinicaltrials.gov: NCT04425629). Non-hospitalized patients > 18 years of age with a SARS-CoV2- positive test result <72 hours, and symptom onset < 10 days before randomization, were randomized 1:1:1 to receive placebo, 2.4g (1.2 g each of casirivimab and imdevimab) or 8.0 g (4.0 g each of casirivimab and imdevimab). The primary virologic endpoint was the time weighted average daily change in viral load from baseline (Day 1) through Day 7. The key secondary clinical endpoint was the proportion of patients with 1 or more Covid-19 related medically-attended visit (MAV) through Day 29. For the present work, 3523 nasopharyngeal, 17 nasal and 6 saliva samples retrieved from a total of 728 patients were analyzed.

##### Hospitalized patient clinical trial

This is an ongoing adaptive, multicenter, randomized, double-blind, placebo-controlled, seamless phase 1/2/3 clinical trial in hospitalized patients with Covid-19 (clinicaltrials.gov: NCT04426695). Hospitalized patients > 18 years of age with a SARS- CoV2-positive test result <72 hours, and symptom onset < 7 days before randomization, were randomized 1:1:1 to receive placebo, 2.4g (1.2 g each of casirivimab and imdevimab) or 8.0 g (4.0 g each of casirivimab and imdevimab). The primary clinical endpoint was the the incidence of death or required mechanical ventilation through Day 29. For the present work, 968 nasopharyngeal, 249 nasal and 119 saliva samples retrieved from a total of 272 patients were analyzed.

#### Virus RNA, RT-PCR and Sequencing

Viral RNA was extracted from hamster samples and from samples collected in the clinical trials (Viracor-Eurofins, MO, USA). RNA*later* Stabilization Solution was aspirated from sample storage tubes, and 1-5mL Qiazol Lysis Reagent (Qiagen, #79306) was added. Tissues were homogenized in Qiazol using an Omni International Prep 96 Homogenizer. The resulting lysate was stored overnight at -20C. In preparation for extraction, lysates were thawed, vortexed, and set up on the deck of a Hamilton Robotics Microlab STAR. The STAR dispensed 150uL of chloroform (Sigma Aldrich, #C2432-500mL) per well to a 2 mL deep well plate (Life Technologies, #4388476), followed by the addition of either 375 or 750uL of lysate. Additional Qiazol was added to wells where only 375uL lysate was added to bring the total volume of lysate to 750uL. Robotic 96-channel pipetting was used to mix the Qiazol and chloroform layers in the deep well plate, followed by a 10 minute incubation at room temperature. The deep well plate was transferred off the STAR to a High Res Biosolutions NanoCell modular robotic system, then centrifuged for 10 minutes at 18G. The plate was returned to the STAR, and using robotic 96-channel pipetting, 400 µL clear aqueous phase was aspirated from the top layer of the plate and transferred to a deep well plate containing 200 µL 100% isopropanol and 10 µL RNA binding beads from the MagMAX-96 for Microarrays Total RNA Isolation Kit (ThermoFisher, #AM1839). The Nanocell system then transferred this plate to a connected KingFisher Flex. Total RNA was purified on the KingFisher Flex using the aforementioned MagMAX kit, according to manufacturer’s specifications. 80uL of Qiagen RNase-Free DNase treatment (Qiagen, *custom order*) was substituted for the MagMAX kit DNase treatment. Once the purification was completed, the elution plate was transferred back to the STAR, and incubated on a Promega MagnaBot 96 Magnetic Separation Device for 5 minutes to remove residual RNA binding beads. The STAR then transferred 2uL aliquots of the purified RNA to plates for the Unchained Labs Lunatic and Advanced Analytical Fragment Analyzer. The Lunatic plate reader was used to obtain concentration and purity data (A_260_/A_280_, A_260_/A_230_), and the Fragment Analyzer was run with a Standard Sensitivity RNA Analysis Kit (Advanced Analytical, #DNF-471-1000) for RNA quality data (RQN, 28s:18s, and DV200 scores). The remaining RNA was transferred using robotic 96-channel pipetting to a FluidX 96-format rack and stored at -80C.

Qualitative and quantitative measurements of viral load were determined by quantitative RT-PCR,10 ul of RNA combined with 25 ng Human Universal Reference RNA (Agilent) was purified by PureBeads (Roche Sequencing). cDNA synthesis was performed using SuperScript™ IV First-Strand Synthesis System (Thermal Fisher) following vendor’s protocol. Then one half of cDNA (10 ul) was used to generate libraries using Swift Normalase™ Amplicon Panel (SNAP) SARS-CoV-2 Panel (Swift Biosciences) following vendor’s protocol. Sequencing was run on NextSeq (Illumina) by multiplexed paired-read run with 2X150 cycles. Overall quality report was generated summarizing the quality of all reads in each sample, along each base pair.

#### Variant calling analysis

Swift amplicon bulk RNA-seq reads were aligned to the SARS-COV-2 reference genome Wuhan-Hu-1 (accession: MN908947) using Minimap2 (v2.17)(Li, 2018). The alignments were sorted by read name, and primers were clipped by the complementary Swiftbiosciences primerclip software (v0.3.8) (https://github.com/swiftbiosciences/primerclip). Duplicate reads were removed using the Picard package (https://github.com/broadinstitute/picard) and target coverage was summarized for each sample with custom scripts. GATK HaplotypeCaller (v4.1.8) (McKenna et al., 2010) was used to perform single nucleotide polymorphism (SNP) calling. At each SNP, the frequency of viral mutations inferred from the sequencing reads were calculated from samtools (v1.9) pileup output(Li et al., 2009) . Variants were flagged if nucleotide positions were covered by at least 10 reads and frequency of mutated reads at these positions was at least 15%.

#### Variant Calling from SARS-CoV-2 Public Genomes

Variant calling from SARS-CoV-2 public genomes was performed as previously described (Baum, 2020). SARS-CoV-2 complete genome sequences were downloaded from GISAID Nucleotide database (https://www.gisaid.org) on February 4th, 2021.

Sequences were curated and genetic diversity of the spike-encoding gene was assessed across high quality genome sequences using custom pipelines. In brief, Blastn was used to align the Wuhan-Hu-1 spike nucleotide sequence (accession: MN908947) against each individual genome. Results were analyzed and presence of the gene was validated if alignment length was greater than 95% with an identity percentage greater than 70%. Homologous spike protein sequences were extracted, translated and aligned to identify amino-acid changes with respect to the reference.

#### Model of RBD-REGN10985-REGN10933-REGN10987

The cryoEM structure of RBD-REGN10933-REGN10987 PDB (6XDG) and the cryoEM structure of RBD-REGN10989-REGN10985 (Fig S4) were superimposed using the software Pymol with the command align for the RBD, where only REGN10933, REGN10987 and REGN10985 are shown.

#### Cryo-EM sample preparation and data collection

Fab fragments of REGN10985 and REGN10989 antibodies were digested with FabALACTICA kit (Genovis) and purified by size-exclusion chromatography. 400 µg of the REGN10985 Fab and 400 µg of REGN10989 Fab were mixed with 200 µg of SARS-CoV-2 RBD and incubated on ice for ∼30 min before being injected into a Superdex 200 increase gel filtration column equilibrated to 50 mM Tris pH 7.5, 150 mM NaCl. Peak fractions containing the REGN10985 Fab - REGN10989 Fab - RBD complex were collected and concentrated using a 10 kDa MWCO centrifugal filter (Amicon). For cryo-EM grid preparation, the protein sample was diluted to ∼1.5 mg/mL and 0.15% PMAL- C8 amphipol was added. 3.5 µL of protein was deposited onto a freshly plasma cleaned UltrAufoil grid (1.2/1.3, 300 mesh). Excess solution was blotted away with filter paper for 4 s at blotting force 0, and plunge frozen into liquid ethane using a Vitrobot Mark IV (Thermo Fisher). The grid was loaded into a Titan Krios (Thermo Fisher) equipped with a K3 detector (Gatan) and 7,763 movies were collected using EPU (Thermo Fisher) at 105,000x magnification, corresponding to a pixel size of 0.85 Å. A dose rate of 15 electrons per pixel per second was used and each movie was 2 seconds, corresponding to a total dose of ∼40 electrons per Å2.

#### Cryo-EM data processing

All cryo-EM data processing was carried out using cryoSPARC v2.14.21(Punjani et al., 2017). The movies were aligned using patch motion correction and patch CTF estimation. An initial set of particles picked using blob picker were subjected to 2D classification to generate templates for template picking. ∼5.5 million particles picked by template picking were subjected to multiple rounds of 2D classification to remove unbound Fabs and particles containing an incomplete complex, resulting in 607,309 ‘good’ complex particles. Ab initio reconstruction with 6 classes generated a single class containing 146,054 particles that corresponded to the full REGN10985 Fab - REGN10989 Fab - RBD complex. Non-uniform refinement of the particles in this class followed by local refinement resulted in a 3.3 Å resolution (FSC=0.143) map that was used for model building. Into this map, we manually placed models of the RBD (taken from PDB code 6M17) and the two Fabs (taken from prior Regeneron antibody structures). These models were then manually rebuilt using Coot and real-space refined against the map using Phenix(Emsley et al., 2010; Liebschner et al., 2019).

### ADDITIONAL RESOURCES

clinicaltrials.gov: NCT04425629 clinicaltrials.gov: NCT04426695

KEY RESOURCES TABLE

## Supplemental Figures

**Figure S1:**
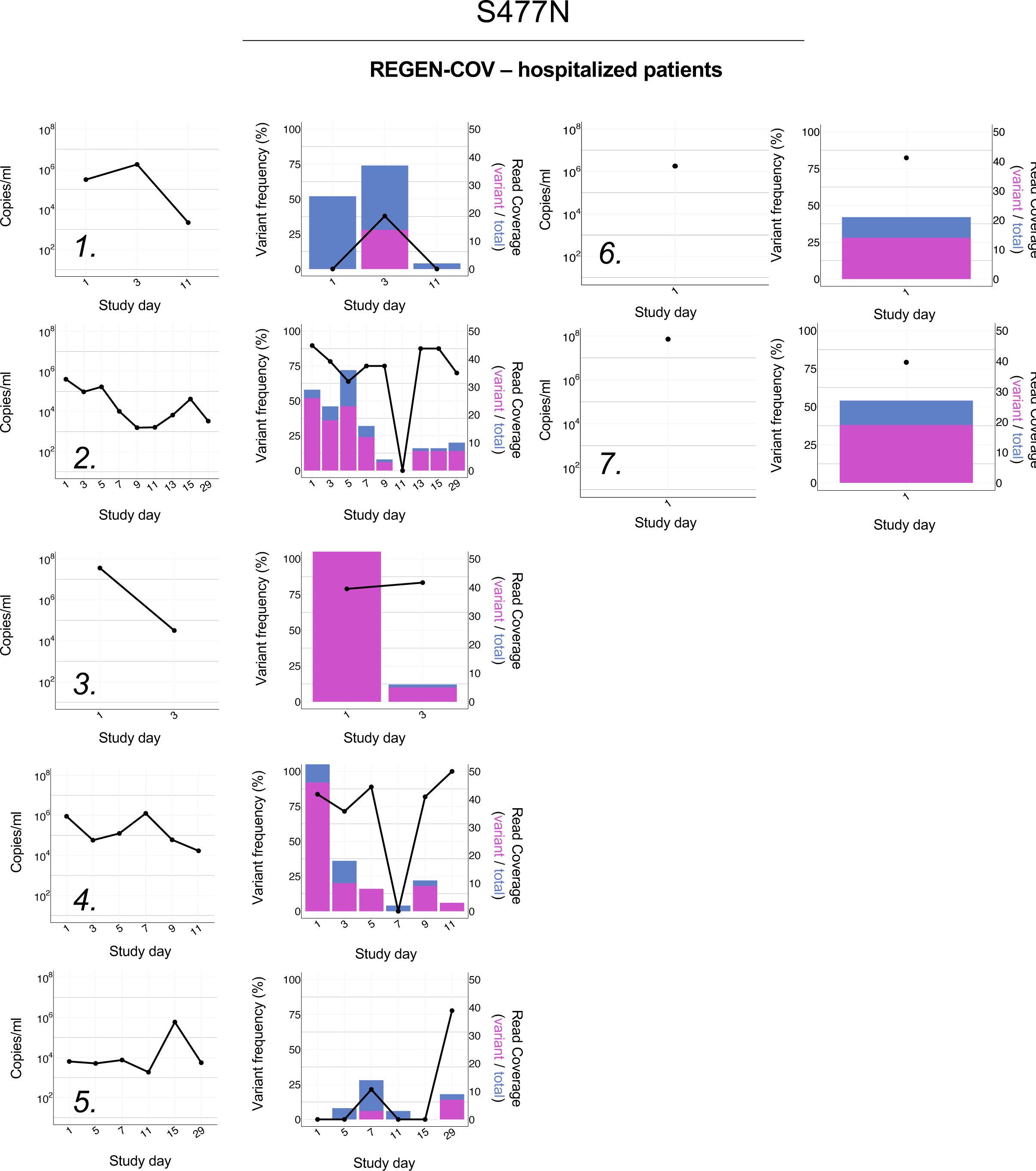
Viral load and longitudinal frequencies of S477N in SARS-CoV-2 positive samples from hospitalized patients. Viral load was determined by quantitative RT-PCR. Variant frequencies are indicated by the black line as percent of reads with nSNPs relative to total number of covering reads. Histograms show total (blue) and mutated (magenta) read coverage at variant position.

**Figure S2:**
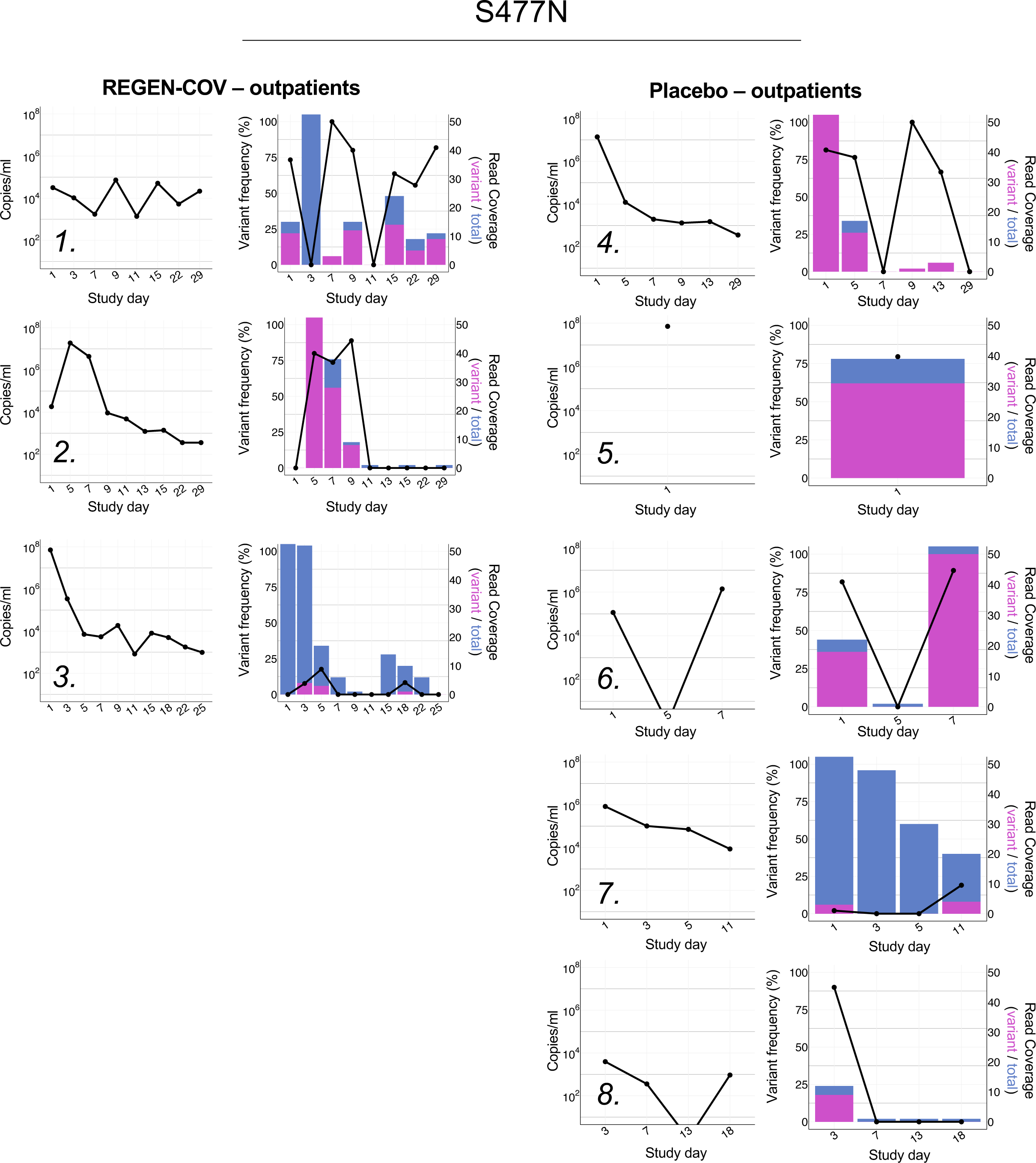
Viral load and longitudinal frequencies of S477N in SARS-CoV-2 positive samples from non-hospitalized patients. Viral load was determined by quantitative RT-PCR. Variant frequencies are indicated by the black line as percent of reads with nSNPs relative to total number of covering reads. Histograms show total (blue) and mutated (magenta) read coverage at variant position.

**Figure S3:**
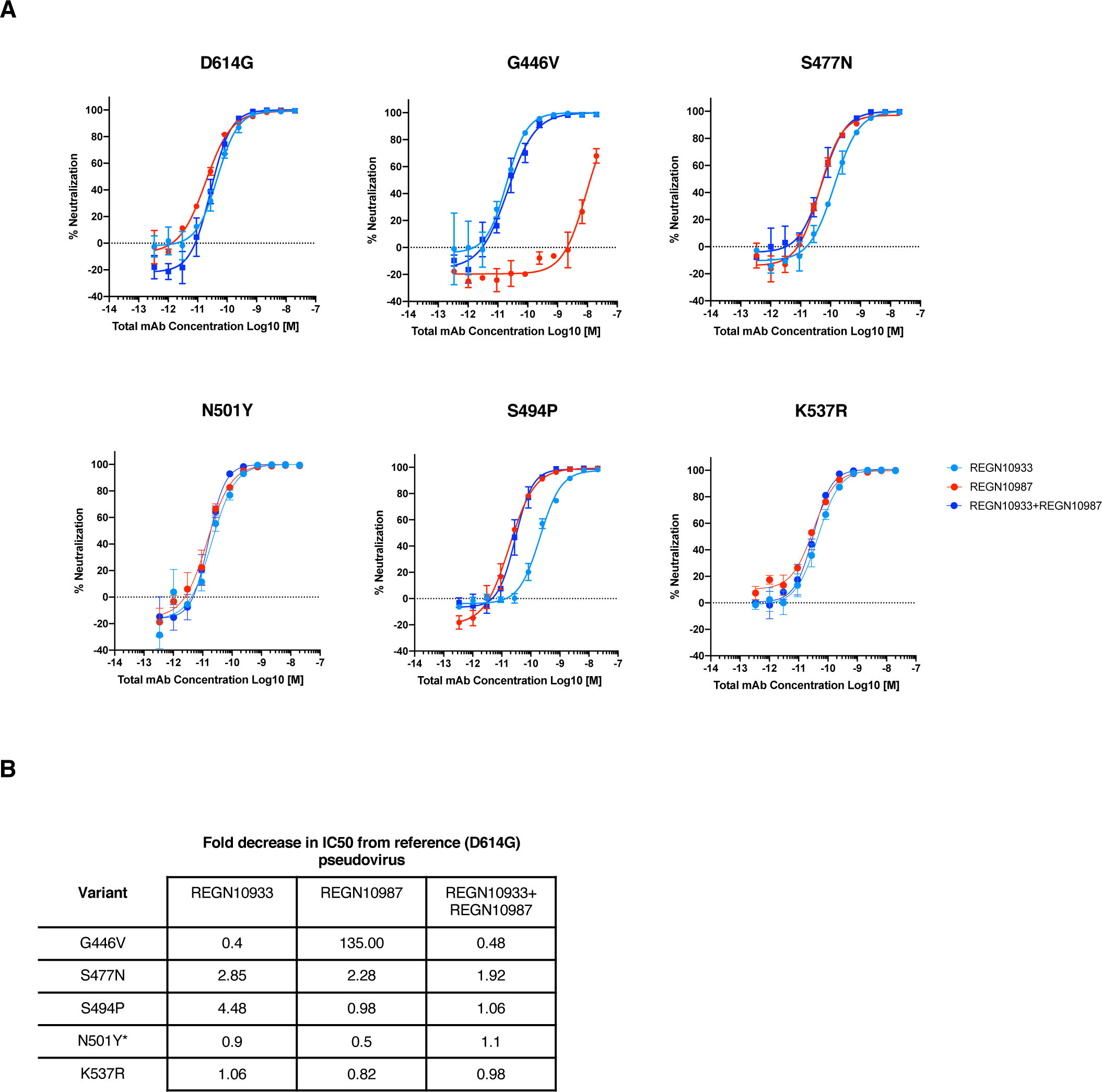
Clinical variant neutralization by the REGEN-COV combination and individual antibodies. Variants identified in two or more samples from at least one patient in either the hospitalized or outpatient trial were assessed for impact on neutralization potency. A) Neutralization curves and B) Fold-change decrease versus parental D614G virus. *The N501Y variant was used to substitute for N501T.

**Figure S4:**
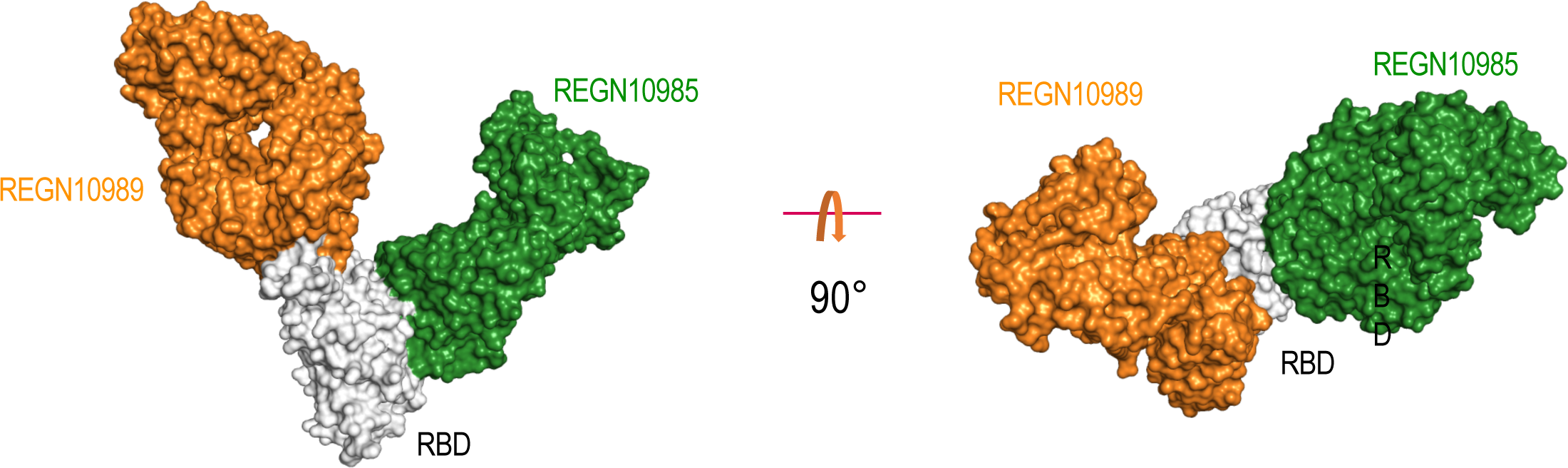
Cryo-EM structure of RBD-REGN10989-REGN10985 in surface representation. RBD is in white, REGN10985 in green and REGN19089 in orange.

**Table S1:** Resistance mutations identified through in vitro escape studies. Mutations is spike protein identified through RNAseq analysis of supernatants from wells with complete escape selected under each antibody condition are listed in the table. The frequency of each mutation from total viral reads at that amino acid position is listed in prentices. Only those mutations that were present at frequencies above 15% are shown as more minor variants are unlikely to contribute to complete escape.

| Antibody | Mutations associated with escape (>15% frequency) | Passage in which escape mutants were identified |
| --- | --- | --- |
| REGN10933 | F486V (88%) | 2 |
| REGN10985 | R408I (40%)<br>G504D (40%) | 1 |
| REGN10987 | K444Q (44%)<br>V445A (40%) | 2 |
| REGN10933+REGN10987 | K417R (100%)<br>K444Q (100%)<br>G476S (90%) | 7 |
| REGN10933+REGN10987+REGN10985 | None | Passaged up to P11 |
| CB6 | K417N (100%) | 1 |
| LY-CoV555 | E484K (38%)<br>Q493R (56%) | 1 |
| CB6+LY-CoV555 | Q493R (100%) | 2 |
| COV2-2130 | K444Q (85%) | 1 |
| COV2-2196 | G476D (81%)<br>N487D (18%) | 2 |
| COV2-2130+COV2-2196 | R346G (98%)<br>E484K (96%)<br>F486V (100%) | 5 |
| VIR-7831 | P337L (16%)<br>T345P (69%) | 2 |

**Table S2:** Summary of variant frequency for each clinical group from study 2066 and 2067. The table includes the patient, sample and amino acid change counts for each study. Frequency of the variants in all samples sequenced is indicated in parenthesis.

**Summary of variant frequency for each clinical group from study 2066 and 2067.**
|  | Study 2066 |  |  | Study 2067 |  |  |
| --- | --- | --- | --- | --- | --- | --- |
|  | Number of patients | Number of samples<br>different time points | Number of variants<br>(variants pers sample) | Number of patients | Number of samples<br>different time points | Number of variants<br>(variants pers sample) |
| **Total** | 272 | 1,336 | 244 (0.18) | 728 | 3,546 | 514(0.14) |
| **Placebo** | 91 | 472 | 90 (0.19) | 243 | 1,215 | 216(0.18) |
| **Treated all** | 181 | 864 | 184 (0.21) | 485 | 2,331 | 399(0.17) |
| **Treated low dose\*** | 88 | 423 | 88 (0.21) | 247 | 1,203 | 248(0.21) |
| **Treated high dose\*** | 93 | 441 | 109 (0.25) | 238 | 1,128 | 212(0.19) |

*\*low dose = 2.4g IV*
*high dose = 8.0g IV*

**Table S3:** Relative neutralization potency of REGEN-COV combination and individual antibodies against spike protein variants identified in hamster lungs. Decrease in neutralization potency relative to the parental virus is shown for each variant identified through RNAseq analysis of hamster lung RNA. * In cases where neutralization potency was not available (N/A) with the exact variant identified in infected hamsters, data is shown for an alternative variant at same position.

|  |  | REGN10933 | REGN10987 | REGN-COV2 |
| --- | --- | --- | --- | --- |
| Variant | Treatment mAb | Decrease in IC50 over reference pseudovirus |  |  |
| P384L | 10987 | 1.91 | 1.29 | 1.65 |
| K444T | 10987 | 1.98 | 1032.52 | 6.42 |
| V445A | 10987 | 1.93 | 548.10 | 4.69 |
| G446S | 10987 | N/A | N/A | N/A |
| G446D | 10987 | N/A | N/A | N/A |
| G446V* | N/A | 0.40 | 134.97 | 0.48 |
| L455W | 10933 | N/A | N/A | N/A |
| L455F* | N/A | 79.54 | 1.06 | 1.79 |
| F486S | 10933 | N/A | N/A | N/A |
| F486L* | N/A | 61.44 | 0.87 | 1.49 |
| F486V* | N/A | 437.54 | 0.87 | 2.68 |

**Table S4:** Sample count and collection days for each clinical group from study 2066 and 2067

| Study 2066 | Total | Placebo | Treated low dose* | Treated high dose* |
| --- | --- | --- | --- | --- |
| Number of Patients | 272 | 91 | 88 | 93 |
| Number of samples different time points | 1336 | 472 | 423 | 441 |
| Baseline | 288 | 95 | 92 | 101 |
| Day2 | 6 | 5 | 1 | 0 |
| Day3 | 223 | 73 | 70 | 80 |
| Day4 | 5 | 2 | 3 | 0 |
| Day5 | 174 | 61 | 60 | 53 |
| Day6 | 5 | 3 | 2 | 0 |
| Day7 | 144 | 53 | 45 | 46 |
| Day8 | 16 | 8 | 4 | 4 |
| Day9 | 119 | 46 | 40 | 33 |
| Day10 | 18 | 8 | 8 | 2 |
| Day11 | 85 | 28 | 18 | 39 |
| Day12 | 16 | 6 | 5 | 5 |
| Day13 | 62 | 30 | 13 | 19 |
| Day14 | 30 | 12 | 6 | 12 |
| Day15 | 51 | 15 | 23 | 13 |
| Day16 | 14 | 7 | 2 | 5 |
| Day17 | 0 | 0 | 0 | 0 |
| Day18 | 1 | 0 | 1 | 0 |
| Day19 | 1 | 0 | 1 | 0 |
| Day20 | 6 | 1 | 2 | 3 |
| Day21 | 3 | 2 | 0 | 1 |
| Day22 | 31 | 9 | 10 | 12 |
| Day23 | 5 | 1 | 2 | 2 |
| Day24 | 1 | 0 | 0 | 1 |
| Day25 | 1 | 0 | 1 | 0 |
| Day26 | 2 | 1 | 1 | 0 |
| Day27 | 1 | 1 | 0 | 0 |
| Day28 | 2 | 0 | 1 | 1 |
| Day29 | 26 | 5 | 12 | 9 |

\*low dose = 2.4g IV high dose = 8.0g IV

| Study 2067 | Total | Placebo | Treated low dose* | Treated high dose* |
| --- | --- | --- | --- | --- |
| Number of Patients | 728 | 243 | 247 | 238 |
| Number of samples different time points | 3546 | 1215 | 1203 | 1128 |
| Baseline | 599 | 205 | 199 | 195 |
| Day2 | 0 | 0 | 0 | 0 |
| Day3 | 534 | 178 | 177 | 179 |
| Day4 | 4 | 0 | 0 | 4 |
| Day5 | 483 | 165 | 163 | 155 |
| Day6 | 29 | 11 | 10 | 8 |
| Day7 | 383 | 120 | 146 | 117 |
| Day8 | 60 | 18 | 19 | 23 |
| Day9 | 301 | 102 | 107 | 92 |
| Day10 | 81 | 26 | 31 | 24 |
| Day11 | 221 | 82 | 73 | 66 |
| Day12 | 73 | 19 | 28 | 26 |
| Day13 | 196 | 78 | 60 | 58 |
| Day14 | 56 | 25 | 13 | 18 |
| Day15 | 150 | 49 | 58 | 43 |
| Day16 | 27 | 8 | 8 | 11 |
| Day17 | 25 | 7 | 13 | 5 |
| Day18 | 82 | 32 | 22 | 28 |
| Day19 | 27 | 9 | 8 | 10 |
| Day20 | 4 | 2 | 2 | 0 |
| Day21 | 26 | 12 | 7 | 7 |
| Day22 | 48 | 17 | 17 | 14 |
| Day23 | 7 | 1 | 1 | 5 |
| Day24 | 15 | 6 | 6 | 3 |
| Day25 | 41 | 20 | 12 | 9 |
| Day26 | 8 | 2 | 3 | 3 |
| Day27 | 8 | 3 | 2 | 3 |
| Day28 | 7 | 4 | 0 | 3 |
| Day29 | 51 | 14 | 18 | 19 |

\*low dose = 2.4g IV high dose = 8.0g IV

